# ER-Anchored Transcription Factors bZIP17 and bZIP60 Regulate Triterpene Saponin Biosynthesis in *Medicago truncatula*^1[OPEN]^

**DOI:** 10.1101/2020.01.17.910802

**Authors:** Bianca Ribeiro, Marie-Laure Erffelinck, Maite Colinas, Clara Williams, Evelien Van Hamme, Elia Lacchini, Rebecca De Clercq, Maria Perassolo, Alain Goossens

**Affiliations:** Ghent University, Department of Plant Biotechnology and Bioinformatics, 9052 Ghent, Belgium; VIB Center for Plant Systems Biology, 9052 Ghent, Belgium; VIB Bio Imaging Core, 9052 Ghent, Belgium; Universidad de Buenos Aires, Facultad de Farmacia y Bioquímica, Departamento de Microbiología, Inmunología y Biotecnología, Cátedra de Biotecnología, Buenos Aires, Argentina; CONICET-Universidad de Buenos Aires, Instituto de Nanobiotecnología (NANOBIOTEC), Buenos Aires, Argentina

## Abstract

Triterpene saponins (TS) are a structurally diverse group of metabolites that are widely distributed in plants. They primarily serve as defense compounds and their production is often triggered by biotic stresses through signaling cascades that are modulated by phytohormones such as the jasmonates (JA). Two JA-modulated basic helix-loop-helix (bHLH) transcription factors (TFs), TRITERPENE SAPONIN BIOSYNTHESIS ACTIVATING REGULATOR 1 (TSAR1) and TSAR2, have been previously identified as direct activators of TS biosynthesis in the model legume *Medicago truncatula*. Here, we report on the involvement of the core endoplasmic reticulum (ER) stress basic leucine zipper (bZIP) TFs bZIP17 and bZIP60 in the regulation of TS biosynthesis. Expression and processing of *M. truncatula* bZIP17 and bZIP60 proteins was altered in roots with perturbed TS biosynthesis or treated with JA. Accordingly, such roots displayed an altered ER network structure. *M. truncatula* bZIP17 and bZIP60 proteins were shown to be capable of interfering with the TSAR-mediated transactivation of TS biosynthesis genes, particularly under ER stress conditions, when they translocate from the ER to the nucleus. Furthermore, the inhibitory role of ER stress bZIP TFs in the regulation of JA-dependent terpene biosynthetic pathways appears to be widespread in the plant kingdom, as we demonstrate that it also occurs in the regulation of monoterpene indole alkaloid biosynthesis in the medicinal plant *Catharanthus roseus*. We postulate that activation of ER stress bZIP TFs provides the plant with a mechanism to balance metabolic activities and thereby adequately govern modulation of growth, development and defense processes in defined stress situations.

**One sentence summary:** ER stress bZIP transcription factors can interfere with the activity of jasmonate-inducible bHLH transcription factors to modulate the elicitation of plant specialized metabolism in stress conditions.

## INTRODUCTION

Plants are continuously challenged with biotic and abiotic stresses. To cope specifically with biotic stresses, such as herbivore feeding or pathogen attack, plants can trigger the biosynthesis of various classes of specialized defense metabolites. A well-known class is that of the structurally and functionally diverse triterpene saponins (TS), which are produced in distinct plant species, including legumes such as *Medicago truncatula*. Phytohormones play an essential role in the stress-induced elicitation of these compounds, again illustrated by the fact that *M. truncatula* TS production is transcriptionally controlled by a JA signaling cascade (Suzuki et al., 2002; De Geyter et al., 2012; Pollier et al., 2013a; Mertens et al., 2016a; Goossens et al., 2017).

TS are amphipathic compounds that consist of lipophilic aglycones (C_30_) covalently linked to one or more hydrophilic sugar moieties (Seki et al., 2015). The diversity of this class of phytochemicals renders them particularly valuable for pharmaceutical and agrochemical purposes (Chang and Keasling, 2006; Ajikumar et al., 2008; Kumari et al., 2013; Moses et al., 2014a; Netala et al., 2015). TS biosynthesis is mainly localized at the ER membrane and shares a biogenic origin with sterols through the mevalonate (MVA) pathway. This pathway, in which 3-HYDROXY-3-METHYLGLUTARYL-COA REDUCTASE (HMGR) is a rate-limiting and heavily regulated enzyme, provides the triterpene precursor isopentenyl pyrophosphate (IPP). Consecutive condensation of six IPP units, catalyzed by prenyltransferases, yields squalene. Squalene is further oxidized by squalene monooxygenase (SQE), generating 2,3-oxidosqualene, which is the last common precursor with the phytosterols. Subsequent cyclization of 2,3-oxidosqualene by saponin-specific 2,3-oxidosqualene cyclases (OSC), more specifically β-AMYRIN SYNTHASE in *M. truncatula*, yields the pentacyclic oleanane-type triterpene backbone β-amyrin (Suzuki et al., 2002; Fukushima et al., 2013; Gholami et al., 2014; Thimmappa et al., 2014; Miettinen et al., 2018; Karunanithi and Zerbe, 2019). Subsequent competitive action of two cytochrome P450-dependent monooxygenases (P450s) results in branching of the *M. truncatula* TS biosynthetic pathway, resulting in the production of two specific classes: the haemolytic and non-haemolytic TS; the former of which were shown to present erythrocyte disruptive activity (Carelli et al., 2011; Gholami et al., 2014; Netala et al., 2015).

The haemolytic TS branch is defined by three consecutive oxidations at position C-28 of β-amyrin by the P450 CYP716A12, thereby yielding oleanolic acid (Carelli et al., 2011; Fukushima et al., 2011). Positions C-2 and C-23 can be further oxidized by respectively CYP72A67 and CYP72A68v2 (Fukushima et al., 2013; Biazzi et al., 2015). The non-haemolytic branch starts with an oxidation reaction at position C-24 of β-amyrin, catalyzed by CYP93E2, and thereby precluding oxidation at position C-28 (Fukushima et al., 2013; Moses et al., 2014b). Subsequent oxidation at position C-22 by CYP72A61v2 yields soyasapogenol B (Fukushima et al., 2013). UDP-dependent glycosyltransferases (UGTs) can further decorate the triterpene aglycones through attachment of sugar moieties, additionally diversifying the TS compendium (Seki et al., 2015).

The past two decades, great progress has been made in the quest for TFs that are modulated by JA or other cues and that control the production of specialized metabolites in plants (De Geyter et al., 2012; Zhou and Memelink, 2016; Goossens et al., 2017; Colinas and Goossens, 2018; Shoji, 2019). Particularly relevant are the basic helix-loop-helix (bHLH) TFs (Goossens et al., 2017). MYC2 was the first bHLH TF reported to control different branches of terpene biosynthesis in *Arabidopsis thaliana*, *Solanum lycopersicum* and *Artemisia annua*, among others (Hong et al., 2012; Kazan and Manners, 2013; Spyropoulou et al., 2014; Goossens et al., 2017). Later, in the medicinal plant *Catharanthus roseus*, source of the anti-cancer drugs vinblastine and vincristine, both MYC2 as well as MYC2-unrelated bHLH TFs, such as BHLH IRIDOID SYNTHESIS 1 (BIS1) and BIS2, were found to elicit the monoterpenoid branch of the monoterpenoid indole alkaloid (MIA) pathway (Zhang et al., 2011; Van Moerkercke et al., 2015; Van Moerkercke et al., 2016; Schweizer et al., 2018). Likewise, the *M. truncatula* orthologs of the BIS TFs, i.e. TRITERPENE SAPONIN BIOSYNTHESIS ACTIVATING REGULATOR 1 (TSAR1) and TSAR2, were reported to transcriptionally regulate the non-haemolytic and haemolytic branch of TS biosynthesis, respectively (Mertens et al., 2016a). TSAR and BIS orthologs were also recently postulated to direct the biosynthesis of antinutritional TS and soyasaponins in *Chenopodium quinoa* and *Glycyrrhiza uralensis*, respectively (Jarvis et al., 2017; Tamura et al., 2018).

Posttranslational regulatory mechanisms of TS biosynthesis have also been described (Hemmerlin, 2013; Erffelinck and Goossens, 2018). Particularly, the JA-inducible really interesting new gene (RING) membrane-anchor (RMA) E3 ubiquitin ligase MAKIBISHI 1 (MKB1) has been reported to control TS biosynthesis in *M. truncatula* by targeting HMGR for degradation by the 26S proteasome (Pollier et al., 2013a). MKB1 forms part of the endoplasmic reticulum (ER)-associated degradation (ERAD) machinery. Generally, this machinery, which is conserved in eukaryotes, monitors the correct folding of membrane and secretory proteins whose biogenesis takes place in the ER. In yeast and mammals, the ERAD machinery also monitors sterol biosynthesis by controlling HMGR stability, albeit through an E3 ubiquitin ligase from another family than the RMA family (Burg and Espenshade, 2011).

When plants are evoked with environmental stresses, a programmed defense response is launched, in which the ERAD, the unfolded protein response (UPR) and other ER stress responses play an important role (Malhotra and Kaufman, 2007; Liu and Howell, 2010b). For instance, ER stress marker genes are often induced during the early stages of immune responses, suggesting that enhanced ER capacity is needed for immunity (Kørner et al., 2015). Eukaryotic cells install signaling networks in response to ER stress, through ER stress sensors that are tethered at the ER membrane. In the model plant *A. thaliana*, several ER stress-specific sensors, including INOSITOL-REQUIRING ENZYME 1 (IRE1), the basic leucine zipper (bZIP) TFs bZIP17 and bZIP28, and the NAC TFs NAC062 and NAC089, have been reported (Liu et al., 2007a; Iwata et al., 2008; Liu and Howell, 2010a; Moreno et al., 2012; Yang et al., 2014a; Yang et al., 2014b; Henriquez-Valencia et al., 2015; Kim et al., 2018). The primary target of IRE1 in response to stress is *bZIP60* mRNA, which is spliced, causing a frame shift, and thereby the elimination of the transmembrane domain of the bZIP60 TF at translation. This truncated version is consequently translocated to the nucleus, where it can install a specific ER stress response. An analogous ER-to-nucleus translocation occurs with bZIP17 and bZIP28, as well during the UPR or other ER stress conditions, but through a different mechanistic process, involving proteolytic cleavage (Liu et al., 2007a, b). The kinase/endoribonuclease IRE1 has already been linked clearly to plant immunity, as it has been reported that loss of function of IRE1 in *A. thaliana* results in a higher sensitivity to pathogen infection and inadequacy to establish systemic acquired resistance (SAR) (Moreno et al., 2012). Similar to the IRE1/bZIP60 branch, the existence of a bZIP17/bZIP28 branch in immunity is suggested, considering their partly overlapping roles with bZIP60 in chemically induced ER stress (Liu et al., 2007a). Conversely, the transcriptional role of bZIP28 and bZIP60 in ER stress responses is antagonized by NONEXPRESSOR OF PR1 GENES 1 (NPR1), a key regulator of salicylic acid (SA)-dependent responses to pathogens (Kørner et al., 2015). This supports a model in which plant immune responses can modulate plant UPR regulation. Despite the growing evidence linking ER stress to immunity, many molecular mechanisms and components involved remain likely elusive.

Here, we explored the regulatory interplay between JA and ER stress signaling in the model legume *M. truncatula*. We demonstrate that *M. truncatula* bZIP17 and bZIP60 can impede transactivation of TS-specific gene promoters by the JA-responsive TSAR1 and TSAR2 and thereby modulate the output of the JA response. We also provide evidence that the interplay between these two responses may be conserved in the plant kingdom by demonstrating that it also occurs in the regulation of terpene biosynthesis in the distinct plant *C. roseus*.

## RESULTS

### Airyscan Imaging Exposes an Altered ER Network Structure in *M. truncatula* MKB1^KD^ Hairy Roots

Previously, we have reported that silencing of *MKB1* in *M. truncatula* hairy roots (MKB1^KD^) results in a dramatic phenotype. In particular, when transferred to liquid medium, MKB1^KD^ hairy roots dissociated into caltrop-like structures (Pollier et al., 2013a). At the metabolite level, it was reported that accumulation of monoglycosylated TS was increased, whereas accumulation of multiple glycosylated TS was decreased in MKB1^KD^ hairy roots compared to control (CTR) hairy roots (Pollier et al., 2013a). Notably, this perturbed TS profile was accompanied by a TS-specific negative transcriptional feedback response in MKB1^KD^ roots (Pollier et al., 2013a). Then, we postulated that this TS-specific negative feedback in MKB1^KD^ hairy roots may be installed to cope with ectopic accumulation of bioactive monoglycosylated TS (Pollier et al., 2013a). Following further reflection on the possible causal link between the perturbation of a part of the ERAD machinery and the perturbation of TS biosynthesis and regulation, we hypothesized that perturbed ER functionality could trigger an ER-inherent mechanism to manage ER capacity and integrity, and thereby (in)directly modulate TS metabolism.

To further investigate this hypothesis, we performed Airyscan imaging to monitor the ER network structure of *M. truncatula* MKB1^KD^ and CTR hairy roots. Hereby, we exploited the fact that the transformed hairy root lines also ectopically express ER-targeted GFP (GFP-KDEL) under the control of a *rolD* promoter, which is used as a visual marker for transformation. The ER network was visually notably altered in MKB1^KD^ hairy roots when compared to CTR lines, for instance exhibiting less of the characteristic three-way junctions in the ER structure (Fig. 1). These phenotypical features were even more pronounced in MKB1^KD^ hairy roots that were elicited with methyl jasmonate (MeJA) compared to mock-treated MKB1^KD^ hairy roots. Interestingly, MeJA treatment of CTR roots also led to a visual alteration of the ER network structure, but the effect was distinct or at least far less pronounced as the effect caused by loss of MKB1 function (Fig. 1).

**Figure 1.**
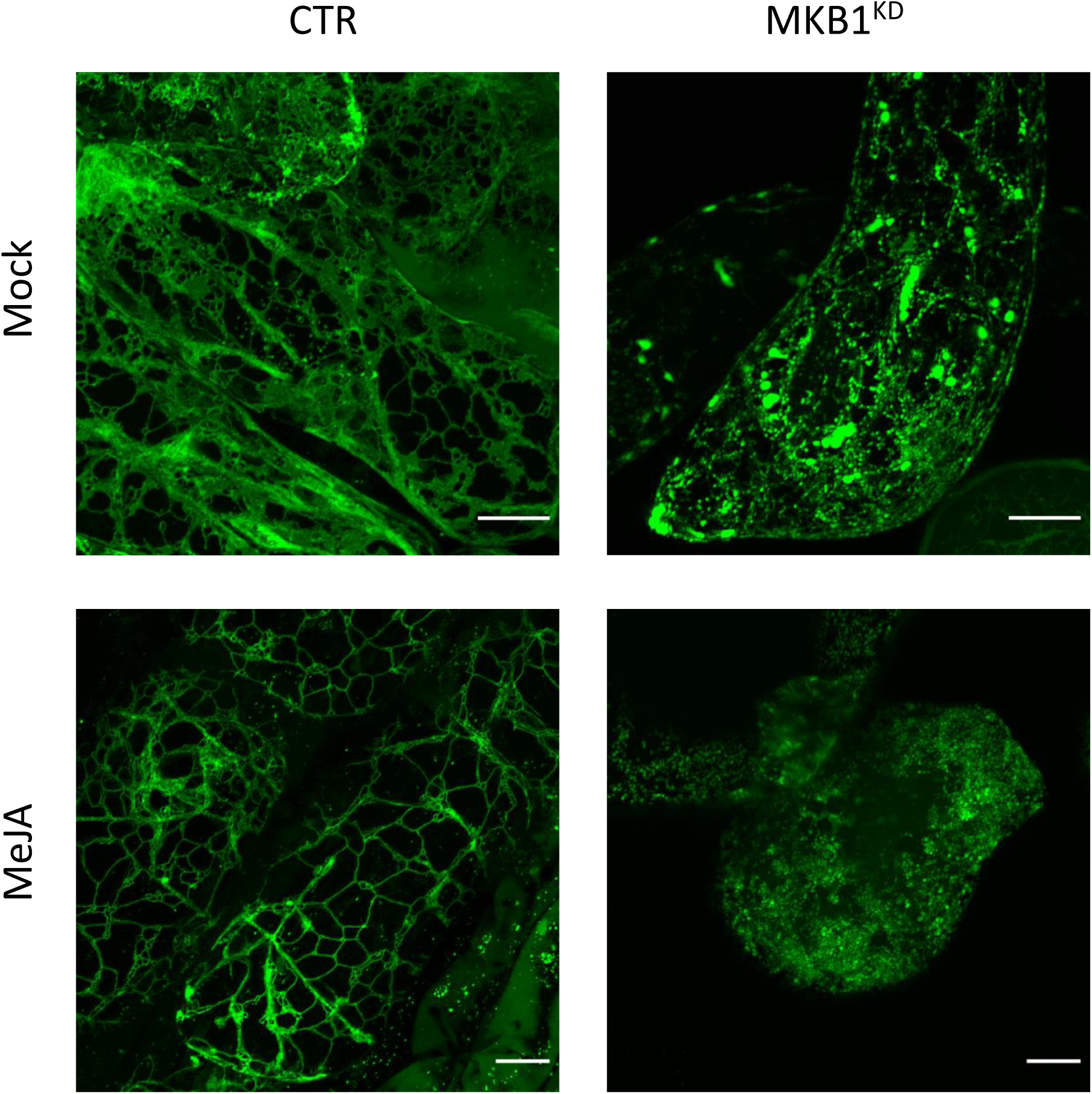
Silencing of *MKB1* and MeJA elicitation alters the ER network structure. Shown are maximum intensity projections of images obtained by Airyscan microscopy of ER-targeted GFP in stably transformed CTR and MKB1^KD^ *M. truncatula* hairy roots. Left, CTR roots elicited (24 h) with ethanol (mock) or 100 µM MeJA (MeJA). Right, mock- and MeJA-treated MKB1^KD^ roots. Scale bars= 20 μm.

### ER Stress Marker Genes Are Transcriptionally Upregulated in MKB1^KD^ Hairy Roots

Given that MKB1^KD^ hairy roots display an altered cell morphology (Pollier et al., 2013a), the differences in ER network structure may not necessarily reflect for instance ER stress caused by the perturbed ERAD machinery but rather an intracellular reorganization following the modifications in the cellular structure. To further assess the possible causes of the altered ER network structure, and/or possibly detect an ER stress response, a transcript profiling study by RNA-sequencing (RNA-Seq) analysis was performed on three independent *M. truncatula* CTR and MKB1^KD^ hairy root lines, either mock- or MeJA-treated. A total of 415,338,234 single-end reads of 50 nt were obtained and mapped on the *M. truncatula* genome version 4.0 (Mt4.0) (Tang et al., 2014). The resulting differential expression profiles were then mined for the closest *M. truncatula* orthologs of a list of known *A. thaliana* ER stress marker genes (Howell, 2013). As such, a set of genes encoding LUMINAL-BINDING PROTEIN 1/2 (BiP1/2; Medtr8g099945) and BiP3 (Medtr9g099795), STROMAL CELL DERIVED FACTOR 2 (SDF2; Medtr3g106130), SORBITOL DEHYDROGENASE (SDH; Medtr1g025430), CALNEXIN (CNX; Medtr3g098430), UDP-glucose:glycoprotein glucosyltransferase (UGGT; Medtr2g006960), HEAT-SHOCK PROTEIN 70 (HSP70; Medtr3g081170), PROTEIN DISULFIDE ISOMERASE-LIKE 1-1 (PDIL1-1; Medtr3g088220), were all found to be significantly upregulated in MKB1^KD^ roots compared to CTR roots, both upon mock- and MeJA treatment (Fig. 2A). Notably, MeJA elicitation itself was also sufficient to trigger an ER stress response in CTR roots, albeit less pronounced (Fig. 2A), in accordance with the moderately altered visual ER network structure (Fig. 1). Together, these data suggest that a transcriptome reminiscent of an ER stress response is not only triggered by loss of MKB1 function but also by JA elicitation.

**Figure 2.**
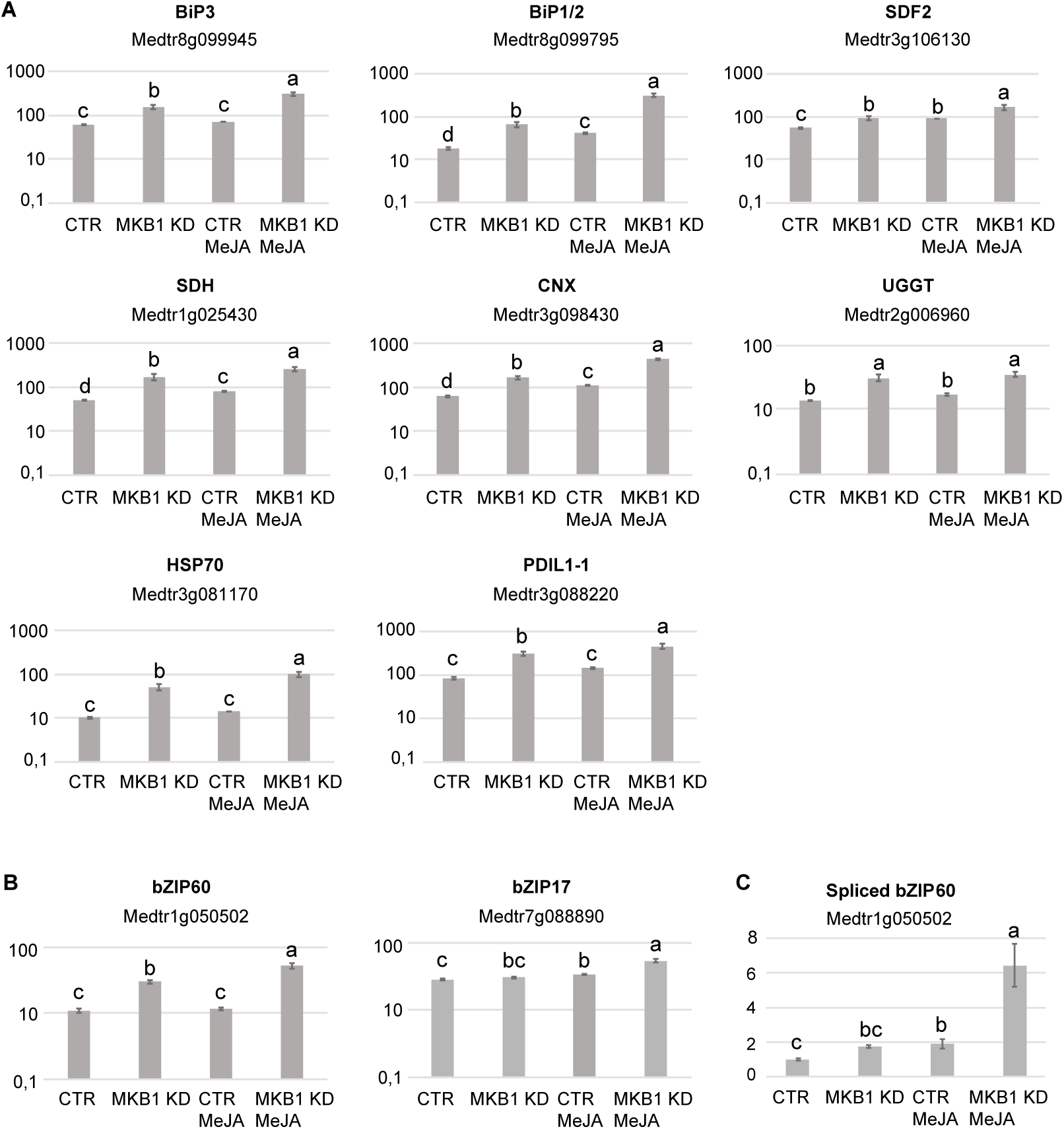
Silencing of *MKB1* and MeJA elicitation both trigger an ER stress response in *M. truncatula*. **A,** RNA-Seq analysis of *M. truncatula* orthologs of *A. thaliana* ER stress marker genes in CTR and MKB1^KD^ roots, mock- or MeJA-treated for 4 h. The Y-axis represents the normalized fragments per kb of exon per million fragments mapped (FPKM) values. Error bars designate SE (n=3, except for MeJA-elicited CTR where n=2). Different letters within each sample indicate statistically significant differences at P < 0.01, as determined by ANOVA, *post hoc* Tukey analysis. **B,** RNA-Seq analysis of *M. truncatula* orthologs of ER stress bZIP TF genes in CTR and MKB1^KD^ roots, mock- or MeJA-treated for 4 h. The Y-axis represents the normalized fragments per kb of exon per million fragments mapped (FPKM) values. Error bars designate SE (n=3, except for MeJA-elicited CTR where n=2). Different letters within each sample indicate statistically significant differences at P < 0.01, as determined by ANOVA, *post hoc* Tukey analysis. **C,** qRT-PCR analysis for detection of spliced *bZIP60* genes in CTR and MKB1^KD^ roots, mock- or MeJA-treated for 4 h. The error bars designate SE (n = 3). Different letters within each sample indicate statistically significant differences at P < 0.01 as determined by ANOVA, *post hoc* Tukey analysis.

Given the above observations, we further assessed the transcript levels of the putative *M. truncatula* orthologs of the core *A. thaliana* ER stress-related basic leucine zipper (bZIP) TFs, *AtbZIP17*, *AtbZIP28* and *AtbZIP60*. Because the bZIP TF gene family is well represented in the *M. truncatula* genome, with at least 81 potential members (Tang et al., 2014), first a phylogenetic analysis for all potential *M. truncatula* bZIP TF genes that were annotated by Wang et al. (2015) was carried out to define the putative *M. truncatula bZIP17*, *bZIP28* and *bZIP60* orthologs. Amino acid sequences of all *M. truncatula* bZIP gene entries, as well as the previously reported *A. thaliana* bZIP TF genes (Dröge-Laser et al., 2018), were retrieved from PLAZA (Van Bel et al., 2018). As previously reported by Dröge-Laser et al. (2018), *AtbZIP17* is part of the group B that also comprises *AtbZIP28* and *AtbZIP49*, whereas *AtbZIP60* is part of the group K as a unique gene. In the *M. truncatula* genome, the bZIP TFs groups B and K are respectively and solely represented by *Medtr7g088890* and *Medtr1g050502* (Supplemental Fig. S1 and Supplemental Fig. S2). It therefore appears that *M. truncatula* might not have other paralogs for the bZIP TF genes in either group B and K, or, alternatively, they are not annotated yet by the Mt4.0 genome, as for instance is also the case for the *MKB1* gene. However, our results are in accordance with a genome-wide analysis of the bZIP TF gene family previously carried out for six legume genomes (*Glycine max, Phaseolus vulgaris, Cicer arietinum, Cajanus cajan, Lotus japonicas*, and *M. truncatula*), where also only a single ortholog for both group B and K bZIP TFs was encountered in five of the legumes studied (Wang et al., 2015). *G. max* formed a notable exception, with two paralogs in each group, which may be the consequence of a recent whole-genome duplication that *G. max* experienced (Wang et al., 2015).

Subsequent mining of the RNA-Seq data indicated that *bZIP17* gene transcript levels were only upregulated in MKB1^KD^ hairy roots elicited with MeJA compared to mock treatment (3.2 fold) (Fig. 2B). No significant difference in *bZIP17* gene transcript levels were observed in MKB1^KD^ roots compared to CTR roots. However, given that activation and translocation of AtbZIP17 occurs posttranslationally following proteolytic cleavage in the Golgi (Liu et al., 2007a; Zhou et al., 2015; Kim et al., 2018), the lack of transcriptional elicitation of *bZIP17* does not exclude that its activity could be enhanced by loss of MKB1 function or by MeJA elicitation. This possibility was not further investigated, however, given that this would demand extensive additional experimentation and that the results for *bZIP60* were more indicative of the activation of an ER stress response. Indeed, *bZIP60* transcripts accumulated to significantly higher levels in MKB1^KD^ hairy roots as compared to CTR roots, both in mock (2.6 fold) and MeJA (3.9 fold) conditions (Fig. 2B). Furthermore, also MeJA treatment could elicit upregulation of *bZIP60* transcript levels particularly in the MKB1^KD^ (2.5 fold) hairy roots (Fig. 2B). Contrary to bZIP17, ‘activation’ of bZIP60 can be assessed at the transcript level, given that it is regulated by IRE1-mediated splicing (Nagashima et al., 2011). To assess the splicing status of *bZIP60*, quantitative reverse transcription PCR (qRT-PCR) was performed using primers designed to detect the predicted spliced *bZIP60* amplicon. This analysis indicated that the level of spliced *bZIP60* amplicons was increased, both in the MKB1^KD^ lines compared to CTR lines and following elicitation with MeJA treatments, both in CTR and MKB1^KD^ hairy roots (Fig. 2C). Together, our transcriptome analysis supports the occurrence of an increased ER stress response, caused by loss of MKB1 function, and to a minor extent also by MeJA elicitation.

### *M. truncatula* bZIP17 and bZIP60 Counteract Transactivation of TS Biosynthesis Promoters by TSAR1 and TSAR2

Because ER stress marker genes are transcriptionally upregulated in MKB1^KD^ hairy roots, we reasoned that this would likely be mediated by the *M. truncatula* bZIP17 and bZIP60 proteins, in analogy with the activity of their putative orthologs in Arabidopsis. We did not investigate this further here, but instead we reasoned that these two bZIP TFs may potentially also negatively regulate TS biosynthesis, hence explaining the TS-specific negative feedback. Therefore, we tested whether bZIP17 and bZIP60 were able to modulate the previously reported transactivation of a set of TS biosynthesis gene promoters by TSAR1 and TSAR2 in a transient expression assay in *Nicotiana tabacum* Bright Yellow-2 (BY-2) protoplasts (Mertens et al., 2016a). More particularly, we used a list of previously generated reporter constructs corresponding to genes encoding enzymes acting at different levels in haemolytic and non-haemolytic TS biosynthesis and consisting of either the 1,000-bp region upstream of the start codon of *HMGR1* (*proHMGR1*), *HMGR4* (*proHMGR4*), *βAS* (*proβAS*), *CYP93E2* (*proCYP93E2*) and *UGT73F3* (*proUGT73F3*), or the 1,500-bp region upstream of the start codon of *CYP72E67* (*proCYP72E67*), all fused to *FIREFLY LUCIFERASE* (*fLUC*) (Mertens et al., 2016a).

First, these reporter constructs were co-transfected in BY2 protoplasts with effector constructs consisting of either the *TSAR1* and *TSAR2* genes driven by the *CaMV35S* promoter. Thereby we could confirm the previously reported transactivation (Mertens et al., 2016a) of *proHMGR1, proHMGR4*, *proβAS* and *proCYP93E2* by TSAR1 (Fig. 3A), and of *proHMGR1, proCYP72E67* and *ProUGT73F3* by TSAR2 (Fig. 3B). Next, we examined the effect of both the full-length *M. truncatula* bZIP17 and bZIP60 and the assumed active and nuclear-localized truncated versions lacking the transmembrane domain, bZIP17Δ and bZIP60Δ, on the transactivation of TS biosynthesis promoters. Therefore, we co-transfected the BY2 protoplasts with the distinct *bZIP* TF effector constructs, which were also driven by the *CaMV35S* promoter. For any of the tested reporter constructs however, no effect of any of the full-length or truncated bZIP TFs on reporter fLUC activity was observed (exemplified with *proCYP93E2* in Supplemental Fig. S3). Finally, we examined the effect of the *M. truncatula* bZIP17 and bZIP60, both the full-length and the truncated versions bZIP17Δ and bZIP60Δ, on the transactivation of TS biosynthesis promoters by TSAR TFs. Remarkably, the high transactivation of *proHMGR1*, *proHMGR4*, *proβAS,* and *proCYP93E2* by *TSAR1*, as compared to the *GUS* control, was significantly repressed when combined with the truncated *bZIP17Δ* or *bZIP60Δ*, but not with intact *bZIP17* or *bZIP60* (Fig. 3A). A similar trend was observed for the transactivation of *proHMGR1*, *proCYP72A67* and *proUGT73F3* by *TSAR2* (Fig. 3B). Taken together, these data suggest that translocated *M. truncatula* ER stress regulatory TFs bZIP17Δ and bZIP60Δ can counteract the transactivation of TS biosynthesis genes in *M. truncatula* by TSAR1 and TSAR2 bHLH factors and may therefore be accountable for the TS-specific transcriptional feedback phenomenon observed in MKB^KD^ hairy roots.

**Figure 3.**
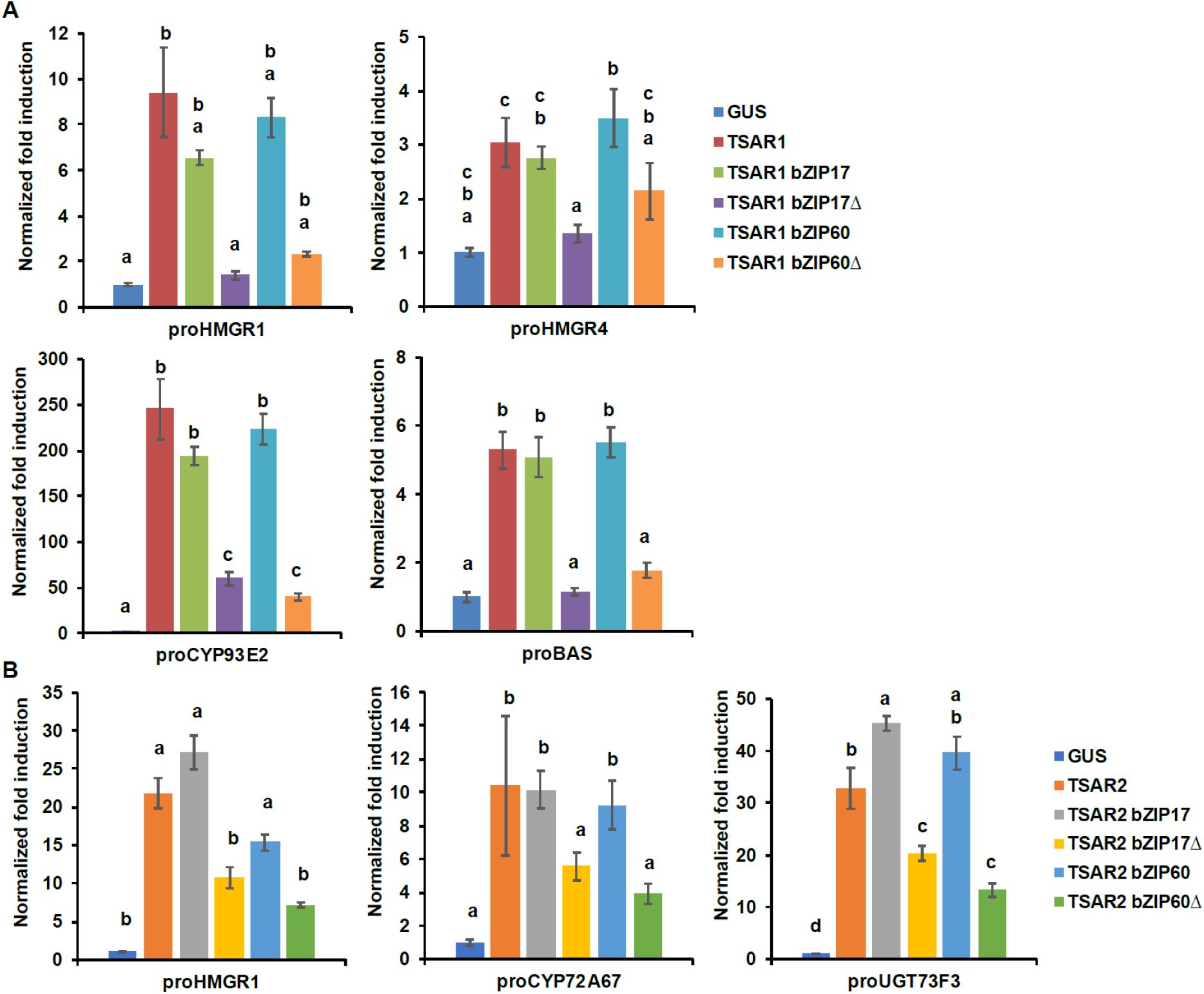
*M. truncatula bZIP17Δ* and *bZIP60Δ* repress transactivation of TS biosynthesis gene promoters by TSAR TFs. **A-B**, Transient transactivation assays in BY-2 protoplasts using the indicated target promoters fused to the *fLUC* reporter gene and TSAR1 (A) or TSAR2 (B) as effectors combined with full-length or truncated bZIP17 and bZIP60. The Y-axis shows fold change in normalized fLUC activity relative to the control transfection with *proCaMV35S:GUS.* The error bars designate SE of the mean (n=8 biological repeats). Different letters within each sample indicate statistically significant differences at P < 0.01, as determined by ANOVA, *post hoc* Tukey analysis.

### What Is the Possible Modus Operandi of bZIP17 and bZIP60 in TSAR Inhibition?

To rationalize the interaction between bZIP17/bZIP60 and TSAR1/TSAR2 we hypothesized two plausible mechanisms wherein induction of TS biosynthesis genes by the bHLH TFs TSAR1/TSAR2 could be antagonized by bZIP17/bZIP60. The first would involve competitive binding of translocated bZIP17/bZIP60 to the TS gene promoters, whereas the second would involve the formation of a protein complex between the translocated bZIPs and the TSARs. In both models, translocated bZIPs would thereby hinder the TSARs in binding or transactivating their target TS biosynthesis promoters.

To assess the first working hypothesis, we first delineated the minimal TS gene promoter sequences needed to observe the bZIP-TSAR1 antagonism. Previously, we demonstrated that the region −101 to −281 relative to the start of the *HMGR1* open reading frame (ORF) (*proHMGR1[−101, −281]*) as well as the region −159 to −300 relative to the start of the ORF of *CYP93E2* (*proCYP93E2[−159,−300]*), are sufficient for transactivation by TSAR1 (Mertens et al., 2016a). Both regions contain one or more of the N-box (5’-CACGAG-3’) TSAR1 target sequences. Using reporter constructs corresponding to these regions in our protoplast assay, bZIP/TSAR1 antagonism is also observed for these minimal *HMGR1* and *CYP93E2* promoter sequences (Fig. 4A-B). In *A. thaliana*, bZIP28 and bZIP60 have been shown to regulate gene expression through binding of ERSE and P-UPRE *cis*-elements in the target gene promoters (Wang et. al 2017). Therefore, we looked for ERSE-I, ERSE-II and UPR sequences in the minimal *proHMGR1[−101,−281]* and *proCYP93E2[−159,−300]* regions. An ERSE-I-like (5’-CCAATATTTTAAGAAGTCAAG -3’) and an ERSE-II-like (5’-ATTCGACCACG-3’) box could be identified in the *proHMGR1[−101,−281]* at position - 211 and *ProCYP93E2[−159,−300]* at position −220, respectively (Fig. 4C-D). To assess whether these ERSE boxes present are necessary for the bZIP-mediated repression of TSAR1 activity, we created mutant promoter versions (*proHMGR1[−101,−281]mut and proCYP93E2[−159,−300]mut)* devoid of these ERSE promoter elements by substitution of the last five bases (5’-CCAATATTTTAAGAAG*TTTTT*-3’ and 5’-ATTCGA*AACAT*-3’) and used the corresponding mutant reporter constructs in transient expression assays in tobacco BY2 protoplasts. Whereas both bZIP17Δ and bZIP60Δ could equally repress TSAR1-mediated transactivation of *proHMGR1[−101,−281]* and *proHMGR1[−101,−281]mut*, repression by bZIP17Δ and bZIP60Δ of TSAR1-mediated transactivation of *proCYP93E2[−159,−300]mut* was abolished or attenuated (Fig. 4A-B), respectively, suggesting that the presence of the ERSE-II box-containing region in *proCYP93E2[−159,−300]* is necessary for bZIP-mediated repression of TSAR1 transactivation. Next, to assess whether bZIP17Δ/bZIP60Δ can directly bind to the ERSE boxes in the TS gene promoters, a yeast one-hybrid (Y1H) assay was performed. To this end, yeast reporter strains harboring integrated synthetic promoters, in which the *proHMGR1* ERSE-I-like box and *proCYP93E2* ERSE-II-like box were repeated in triplicate (3xCCAATATTTTAAGAAGTCAAG [HMGR1] and 3xATTCGACCACG[CYP93E2]), were generated. Unfortunately, binding of bZIP17Δ, bZIP60Δ, TSAR1 or TSAR2 to either the wild-type *proHMGR1* or *proCYP93E2* bait constructs could not be detected (Supplemental Fig. S4).

**Figure 4.**
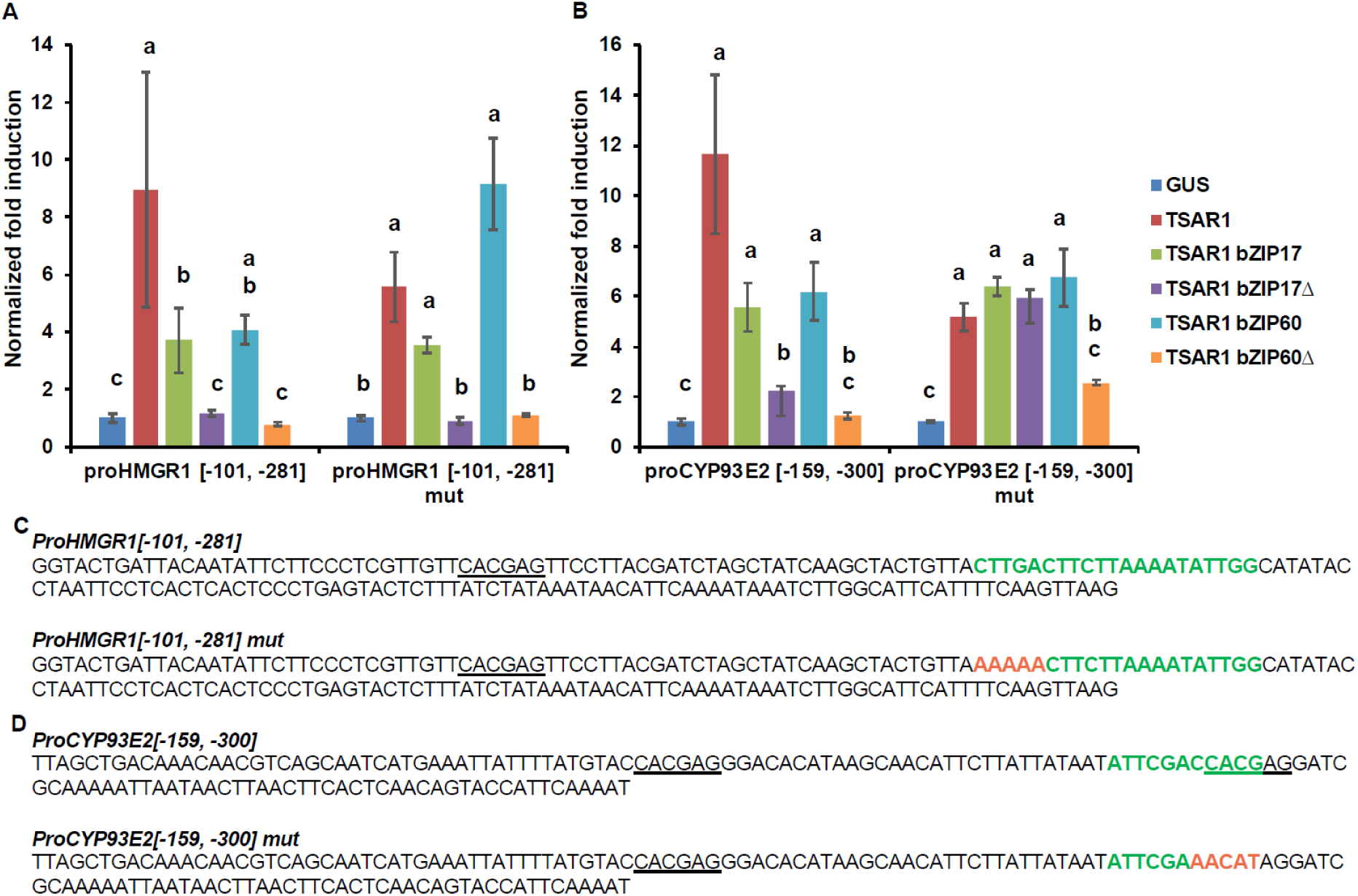
Involvement of ERSE *cis*-elements in the inhibition of TSAR1-mediated transactivation of TS gene promoters by *M. truncatula bZIPs*. **A-B**, Transactivation assays in BY-2 protoplasts using *proHMGR1[−101,−281]* **and** *proHMGR1[−101,−281]mut* (A) or *proCYP93E2[−159,−300]* and *proCYP93E2[−159,−300]mut* (B) as target promoters of different TSAR1 and bZIP17/bZIP60 effector combinations. The Y-axis shows fold change in normalized fLUC activity relative to the control transfection with *proCaMV35S:GUS.* The error bars designate SE of the mean (n=8). Different letters within each sample indicate statistically significant differences at P < 0.01, as determined by ANOVA, *post hoc* Tukey analysis. **C-D,** Sequences of the minimal *HMGR1* and *CYP93E2* promoters. **C,** Nucleotide sequences of *proHMGR1[−101,−281]* and *proHMGR1[−101,−281]mut.* The ERSE-I box (CCAATATTTTAAGAAGTCAAG) is present as reverse complement (CTTGACTTCTTAAAATATTGG) in this sequence. **D,** Nucleotide sequences of *proCYP93E2[−159,−300]* and *proCYP93E2[−159,−300]mut.* Shown is the presence of N- (underlined) and ERSE boxes (in green) in the promoters. The substituted nucleotides in the *proHMGR1[−101,−281]mut* and *proCYP93E2[−159,−300]mut* are shown in orange.

To assess the second working hypothesis, i.e. possible protein complex formation between bZIP17Δ/bZIP60Δ and TSAR1/TSAR2, first a Y2H assay was performed. The outcome of this analysis was negative and did not support possible direct interaction between translocated bZIP17Δ and TSAR1 or TSAR2, at least not in a yeast background (Supplemental Fig. S5). The interaction between bZIP60Δ and TSAR1 or TSAR2 could not assessed by Y2H because of auto-activation of both TSARs and bZIP60Δ. Therefore, we next assessed possible interaction between TSARs and bZIPs through the alternative, *in planta* method, bimolecular fluorescence complementation (BiFC) in agro-infiltrated *Nicotiana benthamiana* leaves. This analysis indicated that at least TSAR1 can interact with both bZIP17Δ and bZIP60Δ (Fig. 5). No interaction could be observed between TSAR2 and either bZIP17Δ and bZIP60Δ (Fig. 5), but, nonetheless, considering all of our data, we postulate that the formation of a protein complex between translocated bZIP TFs and TSAR TFs hinders TSAR-mediated transactivation of TS biosynthesis genes, possibly either by inhibiting DNA binding or recruitment of the RNA transcription machinery.

**Figure 5.**
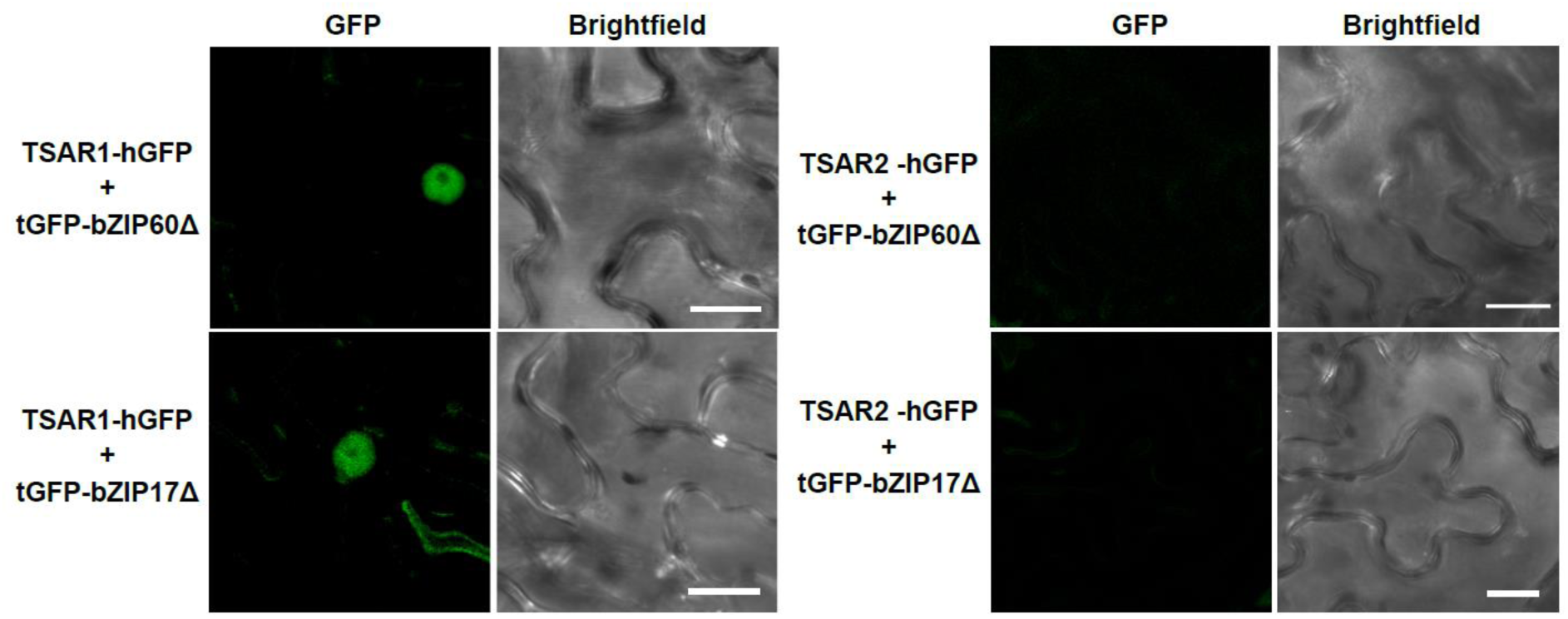
Analysis of physical interaction between bZIP17Δ/bZIP60Δ and TSAR1/TSAR2 by BiFC. Confocal microscopy analysis of *N. benthamiana* leaves agro-infiltrated with constructs expressing fusion proteins of TSAR1 or TSAR2 with a C-terminal fragment of GFP (hGFP) and bZIP17Δ or bZIP60Δ with a N-terminal fragment of GFP (tGFP). Left to right: green, GFP fluorescence; bright field. Scale bars = 20 μm.

### Functional Characterization of *M. truncatula* bZIP17 and bZIP60 *In Planta*

To further evaluate the *in planta* role of the interaction between bZIP17/bZIP60 and TSAR1/TSAR1 in *M. truncatula*, we generated gain- and loss-of-function hairy root lines for the *bZIP17/bZIP60* genes. Three independently generated root lines expressing the *GUS* gene were used as the control. For *bZIP17*, we generated three independent *M. truncatula* hairy root lines overexpressing *bZIP17Δ* (bZIP17Δ-OE, Supplemental Fig. S6A). Quantitative reverse transcription-PCR (qRT-PCR) analysis confirmed overexpression of *bZIP17Δ* by approximately eightfold (Supplemental Fig. S6B). An average fourfold increase of *bZIP60* transcript levels was also observed in those lines. Accordingly, the expression level of the chaperone *BiP1/2* was significantly increased in the bZIP17Δ-OE hairy root lines, in line with previous observations in other species such as *A. thaliana* (Li et al., 2017) and maize (Yang et al., 2013). Although some TS genes did show some differential expression, no consistent significant effect on TS pathway gene expression could be observed in the three bZIP17Δ-OE lines (Supplemental Fig. S6C). Unfortunately, we did not manage to generate lines overexpressing bZIP60Δ, despite several transformation rounds.

We also managed to generate three independent *bZIP17* and two independent *bZIP60* knock-down lines (bZIP17^KD^ and bZIP60^KD^), all showing approximately a fourfold reduction in *bZIP* expression (Fig. 6B and Supplemental Fig. S7B). A notable growth phenotype with a callus-like morphology was observed, especially for bZIP60^KD^ hairy root lines, distinct from the MKB1^KD^ phenotype though (Fig. 6A and Supplemental Fig. S7A). Furthermore, *bZIP60* transcript levels were significantly increased in the bZIP17^KD^ lines (Fig. 6B). Similar observations of a feedback loop to the other ER stress branch were made for the *bZIP60* transcript levels in a double *bzip17 bzip28* mutant in *A. thaliana* line (Kim et al., 2018). Conversely, no feedback on *bZIP17* expression was observed in the bZIP60^KD^ hairy root lines (Supplemental Fig. S7B). Importantly however, the transcript levels of all analyzed TS genes, except for those corresponding to *TSAR1,* were slightly increased in the *bZIP17^KD^* lines (Fig. 6C). In the *bZIP60^KD^* hairy root lines, the effect was less consistent, with only a slight but significant increase in the expression level of *CYP716A12* and *UGT73F3* (Supplemental Fig. S7C). Unfortunately, we did not manage to generate hairy root lines silencing both *bZIP17* and *bZIP60*, despite several transformation attempts. It is plausible to assume that because of the crucial roles of these bZIP factors for plant physiology, simultaneous loss-of-function of both, is not viable. Accordingly, homozygous triple *bzip17 bzip28 bzip60 A. thaliana* mutants present a dwarf phenotype with infertile flowers (Kim et al., 2018). Nonetheless, the data obtained with the *M. truncatula bZIP17^KD^* line support a potential role of at least bZIP17 as a negative attenuator of the TS pathway.

**Figure 6.**
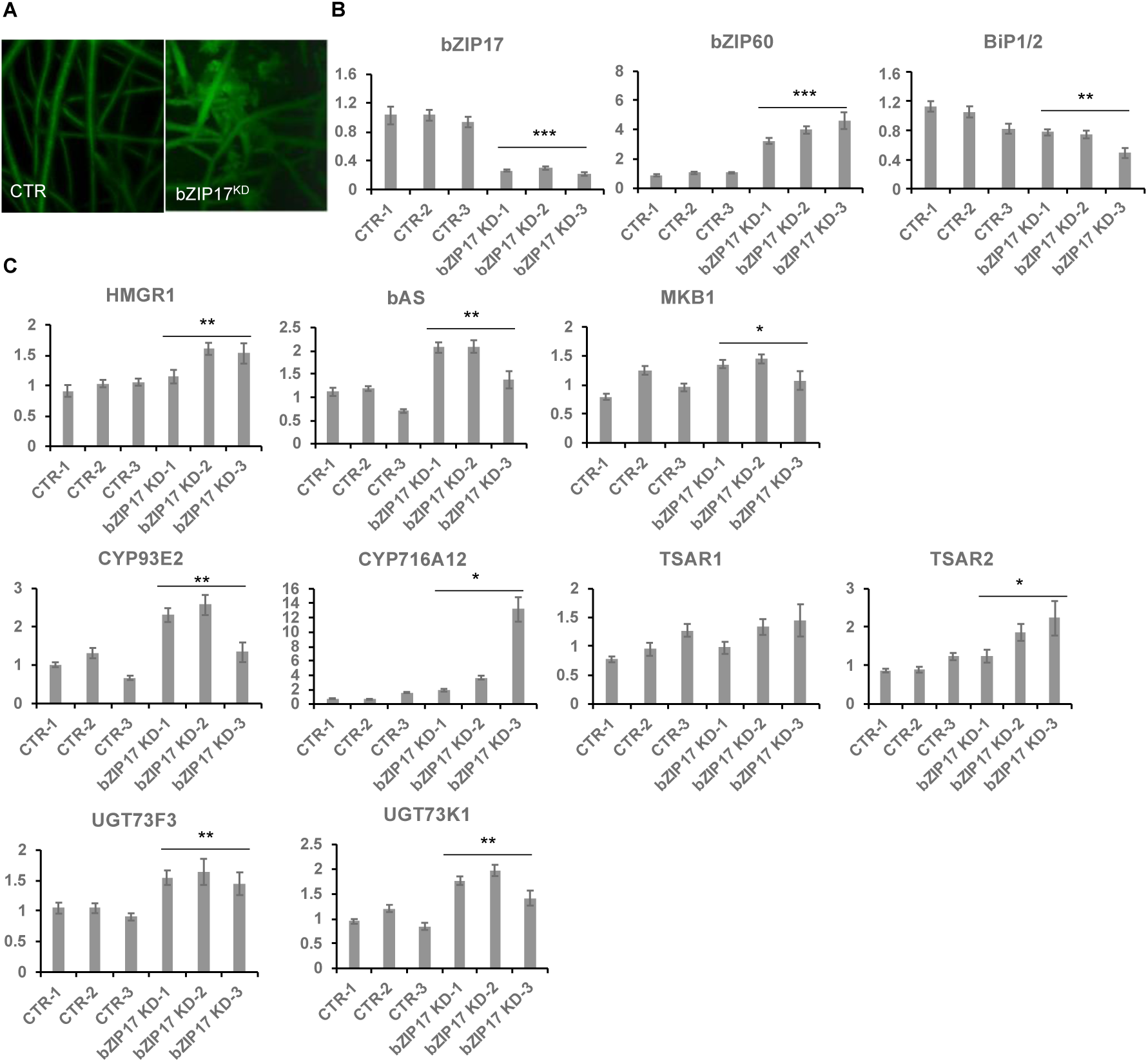
Silencing of *bZIP17* slightly increases TS biosynthesis gene expression in *M. truncatula* hairy roots. **A**, Morphology of control (CTR) and bZIP17^KD^ hairy roots. **B,** qRT-PCR analysis of *bZIP17*, *bZIP60*, *BiP1/2* genes in three independent CTR and *bZIP17^KD^* hairy root lines. **C,** qRT-PCR analysis of TS biosynthetic genes in three independent CTR and *bZIP17^KD^* hairy root lines. Values in the y-axis represent the expression ratio relative to the normalized transcript levels of CTR lines. Three technical replicates were made for each hairy root line. The error bars designate SE (n = 3, technical repeats). Statistical significance was determined by a Student’s *t*-test (*P<0.05, **P<0.01, ***P<0.001).

### ER Stress Inducers Repress Transcript Levels of TS Pathway Genes

Often, TFs that regulate specialized metabolite biosynthetic pathways are coexpressed with the target genes that encode the enzymes of the pathways, particularly when it concerns JA-modulated pathways (Pauwels et al., 2009; De Geyter et al., 2012). In order to find plant growth or stress conditions in which *M. truncatula bZIP17* and *bZIP60* might show an opposite or correlative expression pattern to those of the TS pathway genes, we mined the *M. truncatula* Gene Expression Atlas (MtGEA; http://bioinfo.noble.org/gene-atlas/) (He et al., 2009). We particularly looked for conditions in which increased *bZIP17* and *bZIP60* expression was observed, in combination with a modulated TS gene expression pattern. Such a situation was encountered for *bZIP17* in roots of *M. truncatula* seedlings grown in the presence of 180 mM NaCl (Supplemental Fig. S8). This was not unexpected, given that in *A. thaliana*, salt stress was reported to invoke ER stress, for which the action of bZIP17 is needed for the stress coping mechanism (Li et al., 2017). This observation suggested that *in planta* situations, in which altered bZIP60 and bZIP17 activity may modulate TS gene expression, can indeed be encountered.

As far as we could judge, the MtGEA did not seem to contain transcriptome data of other stress or growth conditions with a pronounced effect on *bZIP17/bZIP60* expression or a reported ER stress effect. However, in *A. thaliana*, other stress agents, such as the reducing agent dithiothreitol (DTT) and the stress hormone salicylic acid (SA), are known to evoke ER stress and the upregulation and activation of *bZIP17* and *bZIP60* (Moreno et al., 2012; Henriquez-Valencia et al., 2015; Li et al., 2017). Hence, we decided to analyze *bZIP,* ER stress and TS pathway gene expression in the presence of NaCl, DTT, SA, all in combination or not with MeJA, in control *M. truncatula* hairy root lines. Given that we anticipated that the MeJA concentration that we typically employ to elicit TS biosynthesis, i.e. 100 µM, may be too strong and possibly mimic antagonistic effects of the ER stress agents, we first determined the minimal MeJA concentration with which TS pathway gene expression could still be induced. qRT-PCR analysis of control *M. truncatula* hairy root lines indicated that at a concentration of 5 µM MeJA, a pronounced and significant induction of TS pathway gene expression could still be observed (Supplemental Fig. S9). This concentration was used for all further experimentation, except for SA, which, as a reported potent JA antagonist of the JA signaling pathway in *A. thaliana* (Van der Does et al., 2013), was still combined with a MeJA concentration of 100 µM.

As expected, *bZIP17* and *bZIP60* transcript levels were increased upon NaCl and DTT treatment in the control *M. truncatula* hairy roots (Supplemental Fig. S10A-B). Upon SA treatment, *bZIP60* transcript levels were increased but not those corresponding to *bZIP17* (Supplemental Fig. S10C). In all cases, increased *BiP1/2* transcript levels were further indicative of successful ER stress induction by all three stress agents. In most cases, combined application with MeJA aggravated the ER stress as reflected by a further increase in the ER stress gene transcript levels (Supplemental Fig. S10). Next, we assessed TS pathway gene expression in all of the lines and different stress treatments. NaCl treatment did not affect the basal expression of TS pathway genes, nor did it interfere with the MeJA elicitation thereof (Fig. 7A). Different and more interesting trends were observed with the DTT and SA treatments. First, SA had a pronounced inhibitory effect on the MeJA induction of all TS pathway genes tested (Fig. 7B). Whether this is partly or entirely mediated by increased bZIP activity cannot be judged at this stage however, given that in *A. thaliana* other TFs such as ORA59 have also been implicated in the SA-mediated suppression of JA signaling (Van der Does et al., 2013). In this regard, the results obtained with the more specific ER stress agent DTT may be considered more indicative. Indeed, also DTT application significantly suppressed the MeJA elicitation of most of the tested TS pathway genes (Fig. 7C). To assess whether the DTT-suppressive effect was specific for MeJA elicitation of TS pathway genes, we also assayed expression of other known JA marker genes by qRT-PCR, all of them corresponding to known early response JA genes involved in the JA amplification loop (Pauwels et al., 2009). Notably, the transcript levels of *JAZ1*, *LOX*, *MYC2a* and *MYC2b* were significantly increased by DTT treatment (Supplemental Fig. S11). Moreover, DTT boosted MeJA elicitation of those JA pathway genes. The latter observations can likely be explained by the reported need for a reducing environment for the JA-Ile–induced Interaction between COI1 and JAZ1 (Yan et al., 2009). As such, overall, our qRT-PCR analysis indicates that the negative effect of DTT on MeJA elicitation is specific for TS pathway genes.

**Figure 7.**
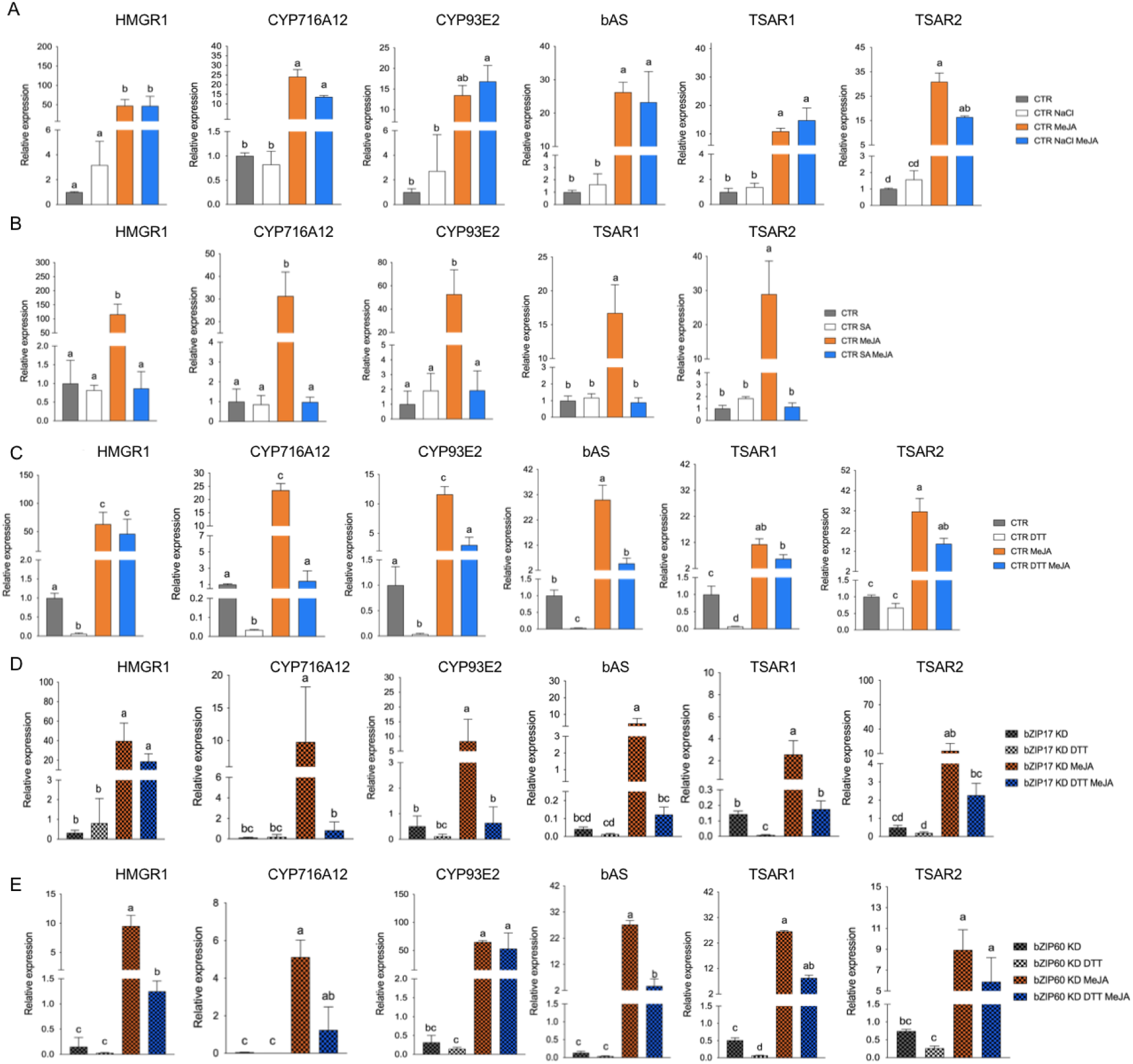
Effects of ER stress on the expression of TS pathway genes in *M. truncatula* hairy roots. **A-E**, qRT-PCR analysis of TS pathway genes in control (CTR) (**A-C**), *bZIP17^KD^* (**D**) and *bZIP60^KD^* (**E**) hairy root lines. Hairy roots were treated with 300 mM NaCl (A), 0.5 mM SA (B) or 2 mM DTT (C-E) for 4 hours prior to the 4-h MeJA treatment (5 µM for A,C-E and 100 µM for B). Expression ratios were plotted relative to the normalized mock-treated line. The error bars designate SE (n = 3 for A-D and n = 2 for E). Different letters within each sample indicate statistically significant differences at P < 0.01, as determined by ANOVA, *post hoc* Tukey analysis.

Finally, we assessed the effect of combined DTT and MeJA application also in the *bZIP17^KD^* and *bZIP60^KD^* hairy root lines. However, no consistent or pronounced differences could be detected with regard to the antagonistic effect that was observed in the control lines (Fig. 7D-E). Only MeJA elicitation of *CYP716A12* and *TSAR2* was less pronounced in both knock-down lines, and the suppressive effect of DTT on MeJA elicitation was only attenuated for *CYP716A12*. We assume that either the MeJA effect is capable of overruling the ER stress, or alternatively that the remaining active *bZIP* gene in the knock-down lines can still account for the ER stress effect. Nonetheless, taken together, our data suggest that also *in planta*, ER stress, and credibly thus the accompanied translocation and activation of bZIP17 and bZIP60, can attenuate the JA response in *M. truncatula* roots, or at least part of it, such as the elicitation of the TS pathway.

### *C. roseus* bZIP17 and bZIP60 Counteract Transactivation of Monoterpenoid Indole Alkaloid Biosynthesis Promoter Genes by BIS1

Since the ER stress response is a conserved mechanism in plants and many specialized metabolite biosynthesis pathways are regulated by TFs that bind G-boxes or closely related boxes, we hypothesized that the action of ER stress bZIP factors on the control of specialized metabolite pathways could be conserved across plant species. An obvious model system to explore this is the medicinal plant *C. roseus*, in which different branches of the monoterpenoid indole alkaloid (MIA) are controlled by bHLH factors such as MYC2 and the BISs, which are functional orthologs of the TSARs (Van Moerkercke et al., 2015; Mertens et al., 2016b; Van Moerkercke et al., 2016; Schweizer et al., 2018).

To determine the putative orthologs of the *bZIP17* and *b*ZIP*60* genes in *C. roseus*, a BLAST analysis was performed using the AtbZIP17 and AtbZIP60 amino acid sequences as query in the *C. roseus* Functional Genomics Database (croFGD; http://bioinformatics.cau.edu.cn/croFGD/) and Medicinal Plant Genomics Resource (http://medicinalplantgenomics.msu.edu/). The highest significant hits were *CROT021933* and *CROT026761*, respectively, which were confirmed to belong to the bZIP TFs group B and K, respectively, by phylogenetic analysis (Supplemental Fig. S12). Moreover, *CROT021933* and *CROT026761* shared the same conserved protein sequence motifs with their bZIP TF group members (Supplemental Fig. S12 and Supplemental Fig. S13).

Next, a transient expression assay was performed in BY2 protoplasts to assess transactivation of a set of promoters from genes encoding enzymes of the *C. roseus* monoterpenoid indole alkaloid (MIA) biosynthesis pathway, including *GERANIOL 8-OXIDASE* (*G8O*), *GERANIOL SYNTHASE* (*GES*) and *IRIDOID SYNTHASE* (*IS*), by the *C. roseus* BIS1 TF in combination with *C. roseus* CrbZIP17, CrbZIP60 or the truncated versions thereof (CrbZIP17Δ and CrbZIP60Δ). And indeed, the transactivation mediated by BIS1 of *proG8O* and *proIS* was compromised in the presence of the truncated *CrbZIP17Δ* and *CrbZIP60Δ* but not of the intact *CrbZIP17* and *CrbZIP60* (Fig. 8A). In case of *proGES*, counteraction of BIS1-mediated transactivation was only observed with *CrbZIP17Δ* (Fig. 8A). Together, these data suggest that translocated ER stress response bZIP TFs can counteract the BIS1-mediated transcriptional activation of MIA biosynthesis genes in *C. roseus*.

**Figure 8.**
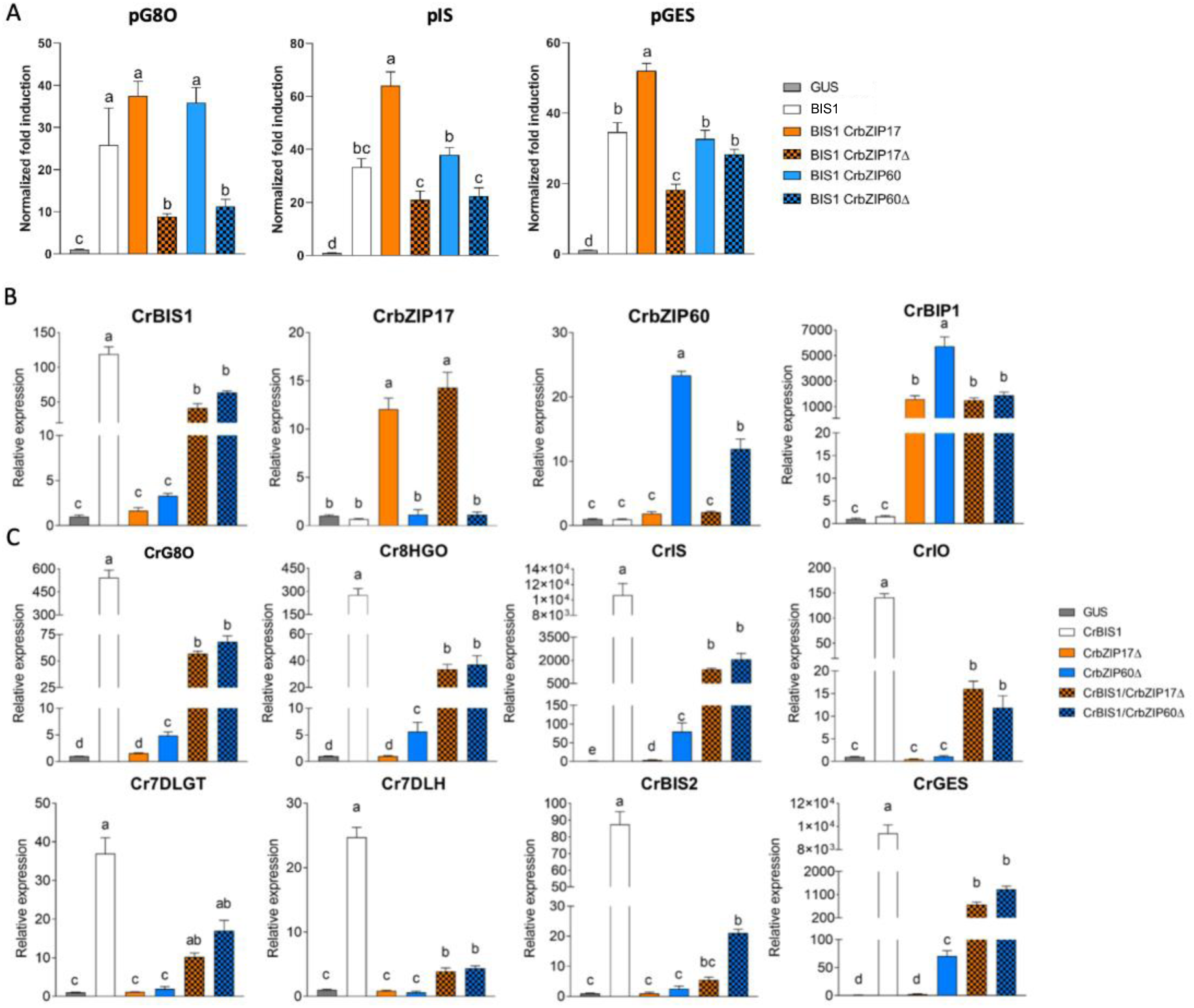
*C. roseus* bZIP17Δ and bZIP60Δ can repress transactivation of MIA biosynthesis by BIS1. **A,** Transient transactivation assays in BY-2 protoplasts using the indicated target promoters fused to the *fLUC* reporter gene and BIS1 as effector combined with full-length or truncated *C. roseus* bZIP17 and bZIP60. The Y-axis shows fold change in normalized fLUC activity relative to the control transfection with *proCaMV35S:GUS.* The error bars designate SE of the mean (n=4). Different letters within each sample indicate statistically significant differences at P<0.01 as determined by ANOVA, *post hoc* Tukey analysis. **B-C,** qRT-PCR analysis of *CrBIS1*, *CrbZIP17*, *CrbZIP60* and *CrBiP1* genes (**B**) and of MIA pathway genes (**C**) in *C. roseus* flower petals transiently overexpressing *CrBIS1*, *CrbZIP17Δ*, *CrbZIP60Δ*, *CrBIS1/CrbZIP17Δ* or *CrBIS1/CrbZIP60Δ* under the *CaMV35S* promoter. Control samples were infiltrated with the *pCaMV35S:GUS* construct. The error bars designate SE of the mean (n=4). Different letters within each sample indicate statistically significant differences at P < 0.01, as determined by ANOVA, *post hoc* Tukey analysis.

To further corroborate this, we also assessed the effect of the *C. roseus* bZIPs in *C. roseus in planta*, in particular through an *Agrobacterium tumefaciens*-assisted *C. roseus* flower infiltration platform that was previously successfully used as an expression system to (co-)express TF(s) and thereby screen for novel MIA biosynthesis regulators (Schweizer et al., 2018). Here, we transiently transformed *C. roseus* flowers with overexpression cassettes for *BIS1*, *CrbZIP17Δ,* and *CrbZIP60Δ*, as well as the double combinations *BIS1/CrbZIP17Δ and BIS1/CrbZIP60Δ.* qRT-PCR analysis confirmed overexpression of all TF genes (Fig. 8B). In support of the functionality of the bZIP TFs, the expression level of the chaperone *BiP1* was significantly increased in all the bZIP17Δ-OE and bZIP60Δ-OE samples (Fig. 8B). Likewise, in line with previous observations (Van Moerkercke et al., 2015; Van Moerkercke et al., 2016; Schweizer et al., 2018), overexpression of *BIS1* strongly upregulated all MIA pathway genes tested (Fig. 8C). Finally, as anticipated, the combinatorial overexpression of *CrbZIP17Δ* or *CrbZIP60Δ* could significantly counteract the BIS1-mediated transcriptional activation of all MIA pathway genes tested (Fig. 8C). Taken together, our findings suggest that interference by ER stress bZIP TFs in the attenuation of JA-dependent terpene biosynthetic pathways might be widespread in the plant kingdom.

## DISCUSSION

The ERAD-type E3 ubiquitin ligase MKB1 has been previously reported to manage TS biosynthesis in *M. truncatula* by controlling HMGR stability (Pollier et al., 2013a). Silencing of *MKB1* in *M. truncatula* hairy roots resulted in an aberrant caltrop-like morphology, increased accumulation of monoglycosylated TS, decreased accumulation of higher glycosylated TS, and a specific downregulation of TS biosynthesis gene expression. Pollier et al. (2013a) speculated that this TS-specific transcriptional feedback might constitute a negative feedback loop to cope with the ectopic accumulation of bioactive monoglycosylated saponins.

Intrigued by this anomaly, we explored the MKB1^KD^ hairy root phenotype by a more in-depth microscopic and transcriptomic analysis. These combined analyses pointed to an ER stress response, reflected by an altered ER network structure and increased transcript levels corresponding to *M. truncatula* orthologs of known *A. thaliana* ER stress marker genes. Because in *A. thaliana*, and conceivably plants in general, the ER stress response can be mediated by two signaling arms, both depending on bZIP TFs, AtbZIP17/AtbZIP28 and AtbZIP60, respectively, that translocate from the ER to the nucleus in ER stress conditions, we speculated that the TS-specific feedback in MKB1^KD^ hairy roots may be mediated by the action of the *M. truncatula* orthologs of these bZIP TFs. Indeed, subsequent functional analysis confirmed that the truncated active versions of *M. truncatula* bZIP60 and bZIP17 could suppress JA and TSAR1/TSAR2-mediated transactivation of TS gene promoters, and could attenuate TS gene expression in *M. truncatula* hairy roots. We therefore postulate that this mechanism could be imposed by plants to regain an equilibrium in ER protein load and folding capacity and/or to fine-tune the biosynthesis of TS under particular stress conditions.

### What Is the Molecular Mode of Action by Which the bZIP TFs Repress TSAR-Induced TS Gene Expression?

Here, we also attempted to determine how the bZIP and TSAR TFs interact to modulate TS biosynthesis. It has been demonstrated in *A. thaliana* that under non-stressed conditions, the activity of bZIP28 in for instance the UPR, which causes ER stress, is inhibited by ELONGATED HYPOCOTYL 5 (HY5), another bZIP TF (Nawkar et al., 2017). This inhibition is mediated by competition for binding to the G-box element (CACGTG) displayed within the ERSE motif(s) in the promoters of the UPR genes. Under ER stress conditions, HY5 undergoes proteasomal degradation, upon which bZIP28 can bind to the ERSE motif(s) and activate the UPR(Nawkar et al., 2017). Conversely, a similar scenario has been reported for JA-inducible genes by Van der Does et al. (2013) in *A. thaliana*, where SA can suppress JA signaling downstream of the JA receptor by targeting GCC promoter motifs via the TF OCTADECANOID-RESPONSIVE ARABIDOPSIS AP2/ERF DOMAIN PROTEIN 59 (ORA 59). In *C. roseus*, the production of MIAs is regulated by the bHLH TF CrMYC2 (Zhang et al., 2015; Paul et al., 2017; Schweizer et al., 2018). Overexpression of *CrMYC2* induces expression of the genes encoding bZIP G-box binding factors (GBFs), resulting in reduced alkaloid accumulation in *C. roseus* hairy roots (Sui et al., 2018). Given that CrGBF1 can bind the same *cis*-element (T/G-box) as CrMYC2 in MIA biosynthesis gene promoters and that CrGBFs can dimerize with CrMYC2, it has been suggested that CrGBF TFs can antagonize CrMYC2 by competitive binding to the T/G-box, and/or by forming a heterodimeric complex, preventing CrMYC2 from binding its target promoters (Sui et al., 2018). Accordingly, we hypothesize a mechanism in *M. truncatula*, in which induction of TS biosynthesis genes by the bHLH TFs TSAR1 and TSAR2 could be antagonized by bZIP17 and bZIP60, either by competitive binding to the promoters or by the formation of a protein complex that would impede TSAR TF transcriptional activity.

ERSE-like *cis-*elements, the targets of the ER stress response bZIPs, are present in several of the TS gene promoters. Notably, in the minimal *CYP93E2* promoter region that contains N-box motifs (5’-CACGAG-3’) that are necessary and sufficient for TSAR-mediated transactivation (Mertens et al., 2016a), this ERSE-like box (5’-ATTCGACCACG-3’) overlaps with one of the N-boxes, making this promoter sequence a plausible target for inhibitory crosstalk. Accordingly, the *M. truncatula* truncated active bZIP17 could no longer suppress the TSAR-mediated transactivation of a mutant version of the *CYP93E2* promoter. It should be noted however that in this mutant promoter version, the partial substitution of the ERSE-like motif also affected the N-box that was part of it, but the other N-box present in the assayed promoter region was left intact. This was also reflected in a reduced capacity of TSAR1 to transactivate this promoter fragment in the absence of truncated bZIPs. Nonetheless, this suggested that competition for binding of *cis*-elements may be a possible mechanism to explain the repressive effect of ER stress bZIP factors on TS gene expression. Conversely however, truncated bZIP60 could still exert its repressive effect on the mutated *CYP93E2* promoter fragment. Likewise, mutations of ERSE-like motifs in another TS gene promoter did not affect the bZIP suppressor effect. Hence, our results are not conclusive and at this stage we can therefore not exclude that there might be other mechanisms at play besides competition for DNA binding to the N-box. A probable alternative mechanism would involve formation of heterodimeric or multimeric protein complexes between bZIPs and TSARs, and/or possibly other proteins involved in transcriptional regulation. Therefore, we also assessed direct binding between TSAR and bZIP TFs using Y2H and BiFC. Again, Y2H was not conclusive, but *in planta* interaction between bZIP17 or bZIP60 and TSAR1 could be established. Plausibly, this interaction hinders TSAR1 transcriptional activity, and, by consequence, transactivation of TS biosynthesis genes is repressed (Fig. 9).

**Figure 9.**
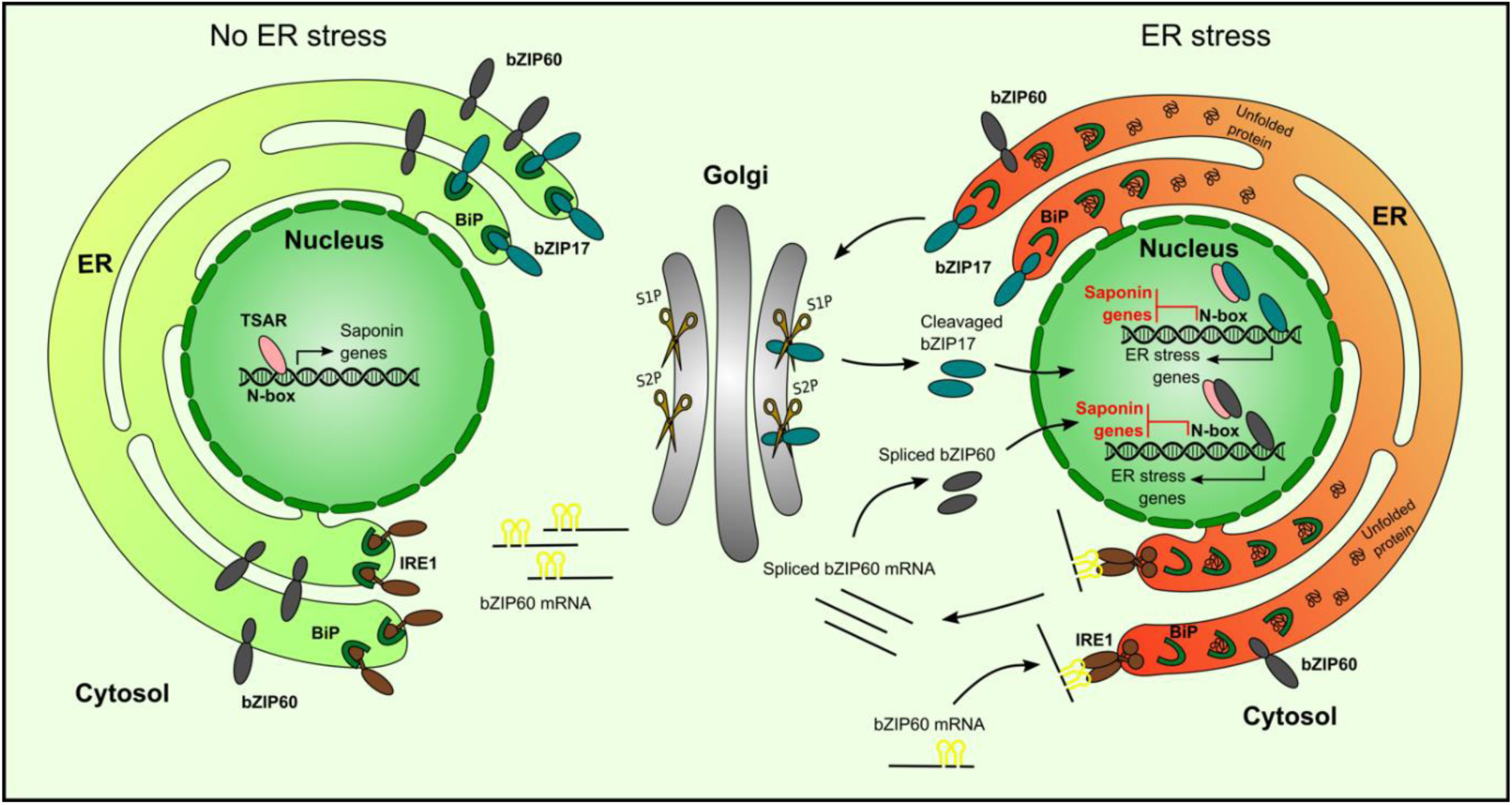
Predicted model of action of bZIP17 and bZIP60 and TSAR under non-ER stress and ER stress conditions. Under non-ER stress conditions, TSAR can bind to the N-box present in the promoters of TS biosynthesis genes and activate their expression. This can occur both in the absence and presence of JAs. Under ER stress conditions, unfolded proteins accumulate and bind to BiP. This triggers IRE1 to perform unconventional splicing of *bZIP60* mRNA as well as release of bZIP17 from the ER and its cleavage by proteases in the Golgi apparatus. The resulting active 30 forms of bZIP17 and bZIP60 translocate to the nucleus, where they can transactivate the expression of ER stress-responsive genes but also bind to the TSARs and thereby repress the transactivation of TS biosynthesis gene expression. Abbreviations: TSAR, triterpene saponin biosynthesis activating regulator; bZIP, basic leucine zipper TF; BiP, binding protein chaperone; IRE1, inositol-requiring enzyme 1; S1P, site-1 protease; S2P, site-2 protease.

### Physiological Relevance and Evolutionary Conservation of the Suppressive Effect of ER Stress-Induced bZIP Factors on JA-Inducible and bHLH TF-Mediated Elicitation of Terpene Biosynthesis in Plants

The molecular mechanisms of ER stress signaling have been well studied over the last years, at least in *A. thaliana*. Many studies made use of compounds such as tunicamycin, which blocks N-linked glycosylation, or the reducing agent DTT, as artificial ER stress inducers. Exploiting the use of such ER stress agents, in particular DTT, we could further support the inverse correlation between TS expression and levels of ER stress, as DTT treatment repressed the expression, particularly in mock conditions and to a lesser extent in the presence of JA, of several TS genes encoding both pathway enzymes and the regulatory TSARs themselves in *M. truncatula* hairy root lines.

There is also increasing knowledge of the physiological conditions upon which ER stress is induced ‘naturally’ in plants. For instance, salt and heat stress has been shown to activate an overall ER stress response in *A. thaliana* (Gao et al., 2008; Deng et al., 2011; Tang et al., 2012; Li et al., 2017). Furthermore, also SA was reported to induce both arms of ER stress signaling in *A. thaliana* (Nagashima et al., 2014), although how is still under debate. It has been reported that SA treatment activates phosphatidylinositol 4-kinase (PI4K), thereby increasing PI phosphates (PIPs) in the Golgi membrane (Krinke et al., 2007). Contrarily, inhibition of PI4K activity leads to a decrease in PIPs and inhibition of *BIP3* induction upon SA treatment (Krinke et al., 2007). Krinke et al. (2007) speculated that upon SA treatment, the phospholipid content in the ER membrane system and traffic within is changed, thereby activating ER-localized ER stress sensors in *A. thaliana*. During SA defense, the transcriptional cofactor NPR1 is converted from oligomers into monomers, which leads to the expression of *PATHOGENESIS-RELATED PROTEIN 1* (*PR1*) genes (Wang et al., 2005). Recently, NPR1 has also been shown to be able to interact with bZIP28 and bZIP60 and to suppress the UPR, independently from SA (Lai et al., 2018). Furthermore, Meng et al. (2017) showed that the CONSTITUTIVE EXPRESSER OF PATHOGENESIS-RELATED GENES 5 (CPR5), a negative modulator of SA, inhibits both the SA-dependent IRE1/bZIP60 and the ER stress-induced bZIP28/IRE1–bZIP60 arms, favoring the growth of plants. Many studies reported both on antagonistic (Rojo et al., 2003; Bostock, 2005; Beckers and Spoel, 2006) and synergistic interactions between SA and JA in plants (Schenk et al., 2000; van Wees et al., 2000. SA appears to confer resistance to biotrophic pathogens, whereas JA to insect herbivory and necrotrophic pathogens {Bostock, 2005 #4; Pieterse et al., 2001; Mur et al., 2006; Stout et al., 2006). This regulatory mechanism could offer the plant a way to activate specific stress response pathways, while repressing others, and, thus contribute to the homeostasis of the cell and/or the appropriate defense response. As indicated above, one mechanism by which SA could antagonize JA action is through the targeting of GCC promoter motifs via the TF ORA59 in *A. thaliana* (Van der Does et al., 2013). Our study suggests that another mechanism of SA–JA antagonism could involve ER stress and the bZIP factors involved therein, but this hypothesis still needs further support with additional experimentation. Nonetheless, such an additional mechanism involving the ER stress machinery could offer plants a means to activate stress response or defense pathways in a stress-specific manner, allowing plants to distinguish between biotic and abiotic stress, or between different pathogens and other attackers. An SA-induced ER stress response could be a mechanism to restrain JA signaling, or at least one of its outputs, namely the elicitation of terpene and/or other specialized metabolite pathways. Given the fact that we observed the antagonism between ER stress-inducible bZIP TFs and JA-inducible bHLH factors in two distinct species, *M. truncatula* and *C. roseus,* which each produce a species-specific compendium of JA-inducible terpene metabolites, this mechanism may be widespread in the plant kingdom. As such, this study may open an avenue for new research on how plants fine-tune their interaction with an ever-changing and often hostile environment.

## MATERIALS AND METHODS

### DNA Constructs

Sequences of the full-length ORFs of *bZIP17* (*Medtr7g088890*) and *bZIP60* (*Medtr1g050502*) were obtained from the *M. truncatula* genome version 4.0 (Tang et al., 2014) and were cloned using Gateway technology (Invitrogen). Full-length and truncated coding sequences for *bZIP17* and *bZIP60* were PCR amplified (for primers, see Supplemental Table S1) and recombined into the donor vector pDONR221. After sequence verification, the entry clones were recombined with the destination vector p2GW7 for *N. tabacum* protoplast assays (Vanden Bossche et al., 2013). The promoter regions of *HMGR1*, *CYP93E2*, *HMGR4*, *CYP72A67, UGT73F3* and *BAS* recombined into the vector pGWL7 (Vanden Bossche et al., 2013) and the full-length coding sequences for *TSAR1* and *TSAR2* recombined with the destination vector p2GW7 had previously been obtained by Mertens et al. (2016a). We generated a fragment of *proHMGR1*, in which the ERSE-I-like 5’-CCAATATTTTAAGAAGTCAAG-3’ box was substituted by 5’-CCAATATTTTAAGAAGTTTTT-3’ by overlap extension PCR. A fragment of *proCYP93E2*, in which the ERSE-II-like 5’-ATTCGACCACG-3’ was substituted with 5’-ATTCGAAACAT-3’, was also constructed by overlap extension PCR. All promoter sequences were recombined into pDONR221, and after sequence verification, entry clones were recombined with the pGWL7 plasmid to generate promoter:fLUC reporter constructs (Vanden Bossche et al., 2013). For the generation of *M. truncatula* hairy roots, sequence-verified entry clones were recombined with the destination vector pK7WG2D for overexpression and pK7GWIWG2(II) for silencing (Karimi et al., 2007). Primers used for cloning of overexpression, silencing in hairy roots and qRT-PCR analysis are reported in Supplemental Table S1.

The coding sequences of *C. roseus* bZIP17 and bZIP60 were amplified from *C. roseus* var. “Little bright eyes” cDNA with Q5® High-Fidelity DNA Polymerase (New England BioLabs®) and recombined into the entry vector pDONR221 (Gateway®). After sequence verification, the entry clones were recombined with the destination vector p2GW7 (Karimi et al., 2002) for *N. tabacum* protoplast assays (Vanden Bossche et al., 2013). The promoter regions of *GES, G10H* and *IS* recombined with the vector pGWL7 (Karimi et al., 2002) and the full-length coding sequence for *BIS1* recombined with the destination vector p2GW7 (Karimi et al., 2002) had previously been obtained by Van Moerkercke et al. (2015). For expression in flower petals under control of the *CaMV35S* promoter, entry clones were recombined into pK7WG2D (Karimi et al., 2002), using LR ClonaseTM enzyme mix (ThermoFisher). The primers used for qRT-PCR analysis are found in Supplemental Table S1. Flower petals of *C. roseus* var. “Little bright eyes” plants (grown under greenhouse conditions) were infiltrated with *A. tumefaciens* C58C1 harboring the constructs for overexpression as previously described (Schweizer et al., 2018).

A Y1H bait fragment of which three identical (CCAATATTTTAAGAAGTCAAG) motifs with their ten flanking nucleotides from the *HMGR1* promoter were fused using two linker sequences was generated by overlap extension PCR. The same was performed with three identical (ATTCGACCACG) motifs with their ten flanking nucleotides from the *CYP93E2* promoter, which were fused using two linker sequences. These constructs were cloned in the reporter plasmid pMW#2 (Deplancke et al., 2006).

### Generation and Cultivation of *M. truncatula* Hairy Roots

Sterilization of *M. truncatula* seeds (ecotype Jemalong J5), transformation of seedlings by *Agrobacterium rhizogenes* (strain LBA 9402/12), and the subsequent generation of hairy roots were carried out as described previously (Pollier et al., 2011). Hairy roots were cultivated for 21 d in liquid medium to provide proper amounts for RNA extraction. For the induction of ER stress, 2 mM DTT was added to the medium for 8 h, and for elicitation of TS pathway gene expression, 100 µM MeJA was added to the medium for 4 h.

### Airyscan Confocal Microscopy

Wild-type and MKB1^KD^ hairy roots were cultivated in nutritive liquid medium (Murashige and Skoog with vitamins supplemented with 1% sucrose) for 2 w and MeJA- or mock-treated for 24 h. Confocal images (16-bit) were captured with an LSM880 confocal microscope equipped with an Airyscan detector (Zeiss, Jena, Germany). Images were taken in super-resolution, FAST mode by using a Plan-Apochromat 63x/1.4 oil objective (1584 × 1584, pixel size: 43 nm × 43 nm). EGFP was excited using the 488 nm line of an Ar laser (30%) and emission was captured between 495 and 550 nm. Z-sections were made every 185 nm. Images were calculated through pixel reassignment and Wiener filtering by using the built-in “Airyscan Processing” command in the Zen software. Fiji was used for generating maximum intensity projections and adding scale bars.

### Phylogenetic Analysis

The bZIP proteins from *A.* and *M. truncatula* were selected based on Dröge-Laser et al. (2018) and Wang et al. (2015), respectively. The amino acid sequences of all selected bZIP proteins were obtained through PLAZA (Van Bel et al., 2018) and the *Catharanthus roseus* Functional Genomics Database (croFGD; http://bioinformatics.cau.edu.cn/croFGD/) (She et al., 2019). These were aligned using MAFFT. The conserved blocks were determined using GBlocks 0.91b and manual curation. IQTREE was used for model selection (Kalyaanamoorthy et al., 2017), after which the best substitution model was selected; the maximum likelihood phylogenetic tree was generated using 1000 bootstrap replicates (Nguyen et al., 2015). The tree figure was made using FigTree software.

### RNA-Seq Analysis

Total RNA of three independent transformant lines per construct was submitted to GATC Biotech (http://www.gatc-biotech.com/) for Illumina HiSeq2500 RNA sequencing (50 nt, single-end read). As described (Pollier et al., 2013b) and using default parameters, the raw RNA-Seq reads were quality-trimmed and mapped on the *M. truncatula* genome v4.0 (Tang et al., 2014) with TOPHAT v2.0.6. Uniquely mapped reads were counted and FPKM values were determined with CUFFLINKS version v2.2.1 (Trapnell, 2013). Differential expression analyses were performed using Cuffdiff (Trapnell, 2013).

### Semi-Quantitative qRT-PCR Analysis

Frozen hairy roots were ground and the material was used to prepare total RNA and first-strand complementary DNA using the RNeasy Mini Kit (Qiagen) and the iScript cDNA Synthesis Kit (Bio-Rad), respectively, according to each manufacturer’s instructions. qRT-PCR primers for *bZIP17* and *bZIP60* were designed using Beacon Designer 4 (Premier Biosoft International) (Supplemental Table S1). The *M. truncatula 40S RIBOSOMAL PROTEIN S8* and *TRANSLATION ELONGATION FACTOR 1A* were used as reference genes. The qRT-PCRs were carried out with a LightCycler 480 (Roche) and the LightCycler 480 SYBR Green I Master Kit (Roche) according to the manufacturer’s guidelines. Three replicates were made for each reaction and the relative expression levels using multiple reference genes were calculated using qBase (Hellemans et al., 2007).

### Transient Expression Assays in Protoplasts

Transient expression assays in *N. tabacum* BY-2 protoplasts were carried out as described by Vanden Bossche et al. (2013). The effector plasmids contained the TSAR, BIS, or bZIP ORFs driven by the *CaMV35S* promoter; the reporter plasmids contained the *fLUC* ORF under control of the target promoters. The normalization plasmid contained the *RENILLA LUCIFERASE* (*rLUC*) under control of the *CaMV35S* promoter. fLUC and rLUC readouts were collected using the Dual-Luciferase® Reporter Assay System (Promega). Each assay incorporated eight biological repeats. Promoter activities were normalized by dividing the fLUC values with the corresponding rLUC values and the average of the normalized fLUC values was calculated and set out relatively to the control fLUC values.

### Y1H Assay

The Y1H reporter strain was obtained as described (Deplancke et al., 2006). *bZIP17* and *bZIP60* full-length and truncated ORFs were cloned into pDEST22 (Invitrogen), to create a fusion with the GAL4 activation domain. *TSAR1* and *TSAR2* recombined with the destination vector pDEST22 (Invitrogen) had previously been obtained by (Mertens et al., 2016a). Empty pDEST22 was used as negative control. The yeast reporter strain was transformed with the preys followed by verification of growth on SD medium lacking His and Trp plates with and without 20 mM 3-amino-1,2,4-triazole after an incubation period of 6 d at 30°C.

### BiFC Assay

*pCaMV35S:ORF-tag* constructs with the N- or C-terminal part of EGFP (nGFP and cGFP, respectively) were constructed by triple Gateway reactions using pK7m34GW (Karimi et al., 2005) as described previously (Boruc et al., 2010). *pCaMV35S:tag-ORF* constructs using the nGFP or cGFP were generated by double Gateway recombination using pk7m24GW2 (Boruc et al., 2010). The constructs were introduced in the *A. tumefaciens* strains C58 for agro-infiltration of the lower epidermal leaf cells of 3- to 4-w-old wild-type *N. benthamiana* plants as described (Boruc et al., 2010). Image acquisition was obtained with a confocal microscope, equipped with a 60x water-corrected objective using the following settings for EGFP detection: EGFP excitation at 488 nm; emission filter 500 - 530 nm. Confocal images were acquired using the ZEN software package attached to the confocal system. Confocal images were processed with ImageJ (www.imagej.nih.gov/ij).

### Y2H Assay

Y2H analysis was performed as described by Cuéllar Pérez et al. (2013) using the GAL4 system, in which bait and prey were fused to the GAL4-AD or GAL4-BD via cloning into pDEST22 or pDEST32, respectively. The *Saccharomyces cerevisiae* PJ69-4α yeast strain was co-transformed with bait and prey constructs using the polyethylene glycol (PEG)/lithium acetate method. Transformants were selected on SD medium lacking Leu and Trp (Clontech, France). Three individual colonies were grown overnight in liquid cultures at 30°C and 10- or 100-fold dilutions were dropped on control (SD-Leu-Trp) or selective (SD-Leu-Trp-His) media. The growth of yeasts was verified after an incubation period of 3 d at 30°C.

## ACKNOWLEDGMENTS

The authors thank Deniz Malat and Robin Vandenbossche for excellent technical assistance, Saskia Lippens for discussions and access to the imaging facilities, and Annick Bleys for the critical reading and help in preparing the manuscript.

